# Molecular diversity analysis of the spike glycoprotein (S) gene from Hong Kong - China

**DOI:** 10.1101/2020.12.16.423166

**Authors:** Eduarda Doralice Alves Braz Da Silva, Dallynne Bárbara Ramos Venâncio, Rosane Maria de Albuquerque, Robson da Silva Ramos, Pierre Teodósio Felix

## Abstract

In this work, 37 haplotypes of spike glycoprotein of SARS-CoV-2 from Hong Kong, China, were used. All sequences were publicly available on the Platform of the National Center for Biotechnology Information (NCBI) and were analyzed for their Molecular Variance (AMOVA), haplotypic diversity, mismatch, demographic and spatial expansion, molecular diversity and time of evolutionary divergence. The results suggested that there was a low diversity among haplotypes, with very low numbers of transitions, transversions, indels-type mutations and with total absence of population expansion perceived in the neutrality tests. The estimators used in this study supported the uniformity among all the results found and confirm the evolutionary conservation of the gene, as well as its protein product, a fact that stimulates the use of therapies based on neutralizing antibodies, such as vaccines based on protein S.

## 1. Introduction

The Spike protein of SARS-CoV-2 (S) is a class I fusion protein, which protrudes on the viral surface for the recognition and binding with the receptor (ACE2) of the host cell, promoting the fusion of the viral membrane with the cell membrane (STERNBERG *et al*., 2020). Once the Spike protein is exposed on the viral surface, it is easily recognized by T cells, which produces a range of neutralizing antibodies from the epitopes and specific domains of S. This protein, which presents a metastable prefusion conformation composed of two functional subunits: the Subunit (S1) responsible for the fusion process still in the viral membrane and the Subunit (S2) responsible for the melting process in the membrane of the host cell, guarantees the entry of the virus into the host cell, having as its only dependence its proteolytic activation, ensuring the process of viral infection only the cleavage of these two subunits (HULSWIT *et al*., 2016).

Some steps of the connection of the virus to the cell surface have been previously described, especially with regard to the proteolysis of the spike protein and the release of its Subunit S2, which is the one that mediates the fusion of the virus and therefore endocytosis. Understanding the critical function of the role of S protein in the virus-host link, has already been serving as a target for therapies with antibodies or chemical compounds, as well as gaining space as a considerable target for vaccines, since serums of mice immunized with SARS-CoV stabilized S protein, significantly reduced the entry of the virus into the target cells, indicating that the cross-neutralizing antibodies, directed to the conserved epitope of S proteins can be produced after vaccination and S-specific neutralizing antibodies, as well as T-cell responses against S protein can be detected within 14 days of vaccination (PILLAY, 2020).

In a historical context, the outbreak and pandemic of *betacoronavirus* SARS-CoV (2002/2003 in China) and MERS-CoV (2012 in Saudi Arabia) gave rise to the development of the first vaccination strategies using recombinant S protein as antigen, due to its high antigenicity and proven ability to induce robust humoral immune responses and neutralizing antibodies in convalescent individuals of SARS-CoV-2 infection (STERNBERG *et al*., 2020). Some other vaccination studies using spike protein in mice and monkeys also induced the formation of S-specific neutralizing antibodies, in addition to protective immunity, strongly evidenced by decreased viral titration in the respiratory tract of animals after contact with SARS-CoV (DU *et al*., 2009).

Other therapeutic approaches, such as the use of phytochemical drugs extracted from Indian medicinal plants, such as *Ocimum sanctum* extract, are also being tried as potent inhibitors of the spike protein for SARS, since it causes a molecular docking in relation to protein-ligant (BASU *et al*., 2020). However, the transformation of peptides and small non-peptide molecules (which target the functional domain of protein S in SARS-CoV, particularly RBD in subunit S1 and HR2 region in subunit S2), into effective and safe antiviral drugs for the treatment of SARS need a significant technological increase in addition to studies that test the in vivo efficiency of these antiviral agents in animal models (DU *et al*., 2009).

Thus, studies on the molecular diversity of the spike protein are increasingly necessary, since it may have different genetic conformations between viral populations, and may have different levels of susceptibility and evasion to immune responses. With this in mind, the team from the Laboratory of Population Genetics and Computational Evolutionary Biology (LaBECom-UNIVISA) conducted a molecular variance study in 37 haplotypes of the Spike protein of SARS-CoV-2, available at the National Center of Biotechnology Information (NCBI).

## 2. Objective

Test the existing molecular variance levels in 37 haplotypes of the spike glycoprotein (S) gene from Hong Kong - China

## 3. Methodology

### Database

The 37 haplotypes of the SARS-CoV-2 spike protein were redeemed from the National Biotechnology Information Center (NCBI) platform and are publicly available at the address (https://www.ncbi.nlm.nih.gov/popset/?term=89474483), accessed December 12, 2020.

### Genetic Structuring Analyses

Molecular Variance (AMOVA), Genetic Distance, mismatch, demographic and spatial expansion analyses, molecular diversity and evolutionary divergence time were obtained with the Software Arlequin v. 3.5 (EXCOFFIER et al., 2005) using 1000 random permutations (NEI and KUMAR, 2000).

All steps of this process are described in: https://dx.doi.org/10.17504/protocols.io.bmbvk2n6 (Felix *et al*., 2020).

## 4. Results

### General properties of analyzed sequences

The 37 sequences analyzed revealed a low level of haplotypic diversity with very low numbers of transitions, transversions and indels-type mutations, with only 17 polymorphic sites (table 1 and table 2 and figure 2).

**Table 1.** Standard molecular diversity index for the 37 SARS-CoV-2 spike protein haplotypes

|  |  |
| --- | --- |
| No. of sequences | 37 |
| No. of usable loci: | 3768 loci (with less than 5.00 % missing data) |
| No. of gene copies: | 37 |
| No. of loci: | 3768 |
| No. of polymorphic sites: | 17 |

**Table 2.** Molecular diversity index for the 37 haplotypes of SARS-CoV-2 SPIKE protein

|  |  |
| --- | --- |
| Sample size | 37 |
| Deletion weight | 1 |
| Transition weight | 1 |
| Transversion weight | 1 |
| Allowed level of missing data | 5% |
| Number of observed transitions | 11 |
| Number of observed transversions | 6 |
| Number of substitutions | 17 |
| Number of observed indels | 0 |
| Number of polymorphic sites | 17 |
| Number of observed sites with transitions | 11 |
| Number of observed sites with transversions | 6 |
| Number of observed sites with substitutions | 17 |
| Number of observed sites with indels | 0 |
| Nucleotide composition (Relative values) |  |
| **C** | 20.05% |
| **T** | 33.30% |
| **A** | 27.97% |
| **G** | 18.69% |

### Molecular Diversity Analyses

The molecular diversity indices estimated for θ indicate that there are no significant mutations between the haplotypes studied, both at the level of transitions and at the level of transversions and indels mutations (insertions and deletions), were also not significant (Table 3). The Tajima D neutrality tests and Fs de Fu, show that there are no disagreements between the general φ and π estimates and their negative and highly significant values, indicating no population expansion.

**Table 3.** Molecular diversity indices for θ values among the 37 haplotypes of the SARS-CoV-2 SPIKE protein

| Statistics | SPIKE | Mean | s.d. |
| --- | --- | --- | --- |
| Theta_k | N.A. | N.A. | N.A. |
| Theta_k_lower | N.A. | N.A. | N.A. |
| Theta_k_upper | N.A. | N.A. | N.A. |
| Theta_H | N.A. | N.A. | N.A. |
| s.d. Theta_H | N.A. | N.A. | N.A. |
| Theta_S | 4.07229 | 4.07229 | 0.00000 |
| s.d. Theta_S | 1.51696 | 1.51696 | 0.00000 |
| Theta_pi | 2.42643 | 2.42643 | 0.00000 |
| s.d. Theta_pi | 1.49590 | 1.49590 | 0.00000 |

The irregularity index (R= Raggedness) with parametric bootstrap simulated new values for before and after a supposed demographic expansion and, in this case, assumed a value equal to zero for the whole group (table 4, table 5 and figure 1).

**Table 4.**
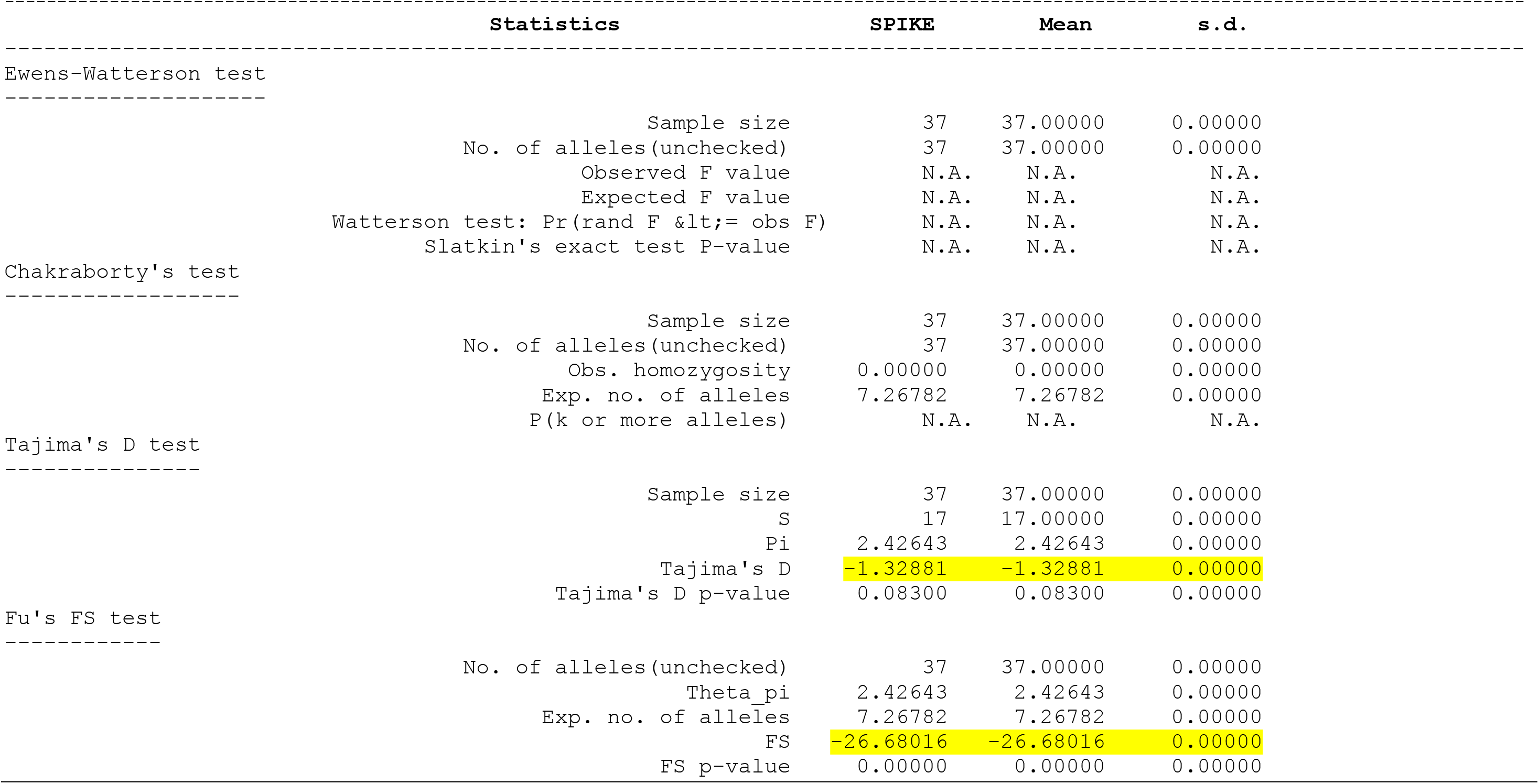
Neutrality test for the 37 haplotypes of the SARS-CoV-2 SPIKE protein

|  | Statistics | SPIKE | Mean | s.d. |
| --- | --- | --- | --- | --- |
| Ewens-Watterson test |  |  |  |  |
|  | Sample size | 37 | 37.00000 | 0.00000 |
|  | No. of alleles(unchecked) | 37 | 37.00000 | 0.00000 |
|  | Observed F value | N.A. | N.A. | N.A. |
|  | Expected F value | N.A. | N.A. | N.A. |
| | Watterson test: $\Pr(\text{rand F} \leq \text{obs F})$ | N.A. | N.A. | N.A. |
|  | Slatkin's exact test P-value | N.A. | N.A. | N.A. |
| Chakraborty's test |  |  |  |  |
|  | Sample size | 37 | 37.00000 | 0.00000 |
|  | No. of alleles(unchecked) | 37 | 37.00000 | 0.00000 |
|  | Obs. homozygosity | 0.00000 | 0.00000 | 0.00000 |
|  | Exp. no. of alleles | 7.26782 | 7.26782 | 0.00000 |
|  | P(k or more alleles) | N.A. | N.A. | N.A. |
| Tajima's D test |  |  |  |  |
|  | Sample size | 37 | 37.00000 | 0.00000 |
|  | S | 17 | 17.00000 | 0.00000 |
|  | Pi | 2.42643 | 2.42643 | 0.00000 |
|  | Tajima's D | -1.32881 | -1.32881 | 0.00000 |
|  | Tajima's D p-value | 0.08300 | 0.08300 | 0.00000 |
| Fu's FS test |  |  |  |  |
|  | No. of alleles(unchecked) | 37 | 37.00000 | 0.00000 |
|  | Theta_pi | 2.42643 | 2.42643 | 0.00000 |
|  | Exp. no. of alleles | 7.26782 | 7.26782 | 0.00000 |
|  | FS | -26.68016 | -26.68016 | 0.00000 |
|  | FS p-value | 0.00000 | 0.00000 | 0.00000 |

**Table 5.** Mismatch analysis: demographic and spatial expansion rates for the 37 haplotypes of the SARS-CoV-2 SPIKE protein

| Statistics | SPIKE | Mean | s.d. |
| --- | --- | --- | --- |
| Demographic expansion |  |  |  |
| Tau | 2.50000 | 2.50000 | 0.00000 |
| Theta0 | 0.00000 | 0.00000 | 0.00000 |
| Theta1 | 6834.95675 | 6834.95675 | 0.00000 |
| SSD | 0.00184 | 0.00184 | 0.00000 |
| Raggedness index | 0.04228 | 0.04228 | 0.00000 |
| Spatial expansion |  |  |  |
| Tau | 2.23800 | 2.23800 | 0.00000 |
| Theta | 0.48500 | 0.48500 | 0.00000 |
| M | 10007.06061 | 10007.06061 | 0.00000 |
| SSD | 0.00362 | 0.00362 | 0.00000 |
| Raggedness index | 0.04228 | 0.04228 | 0.00000 |

**Figure 1.**
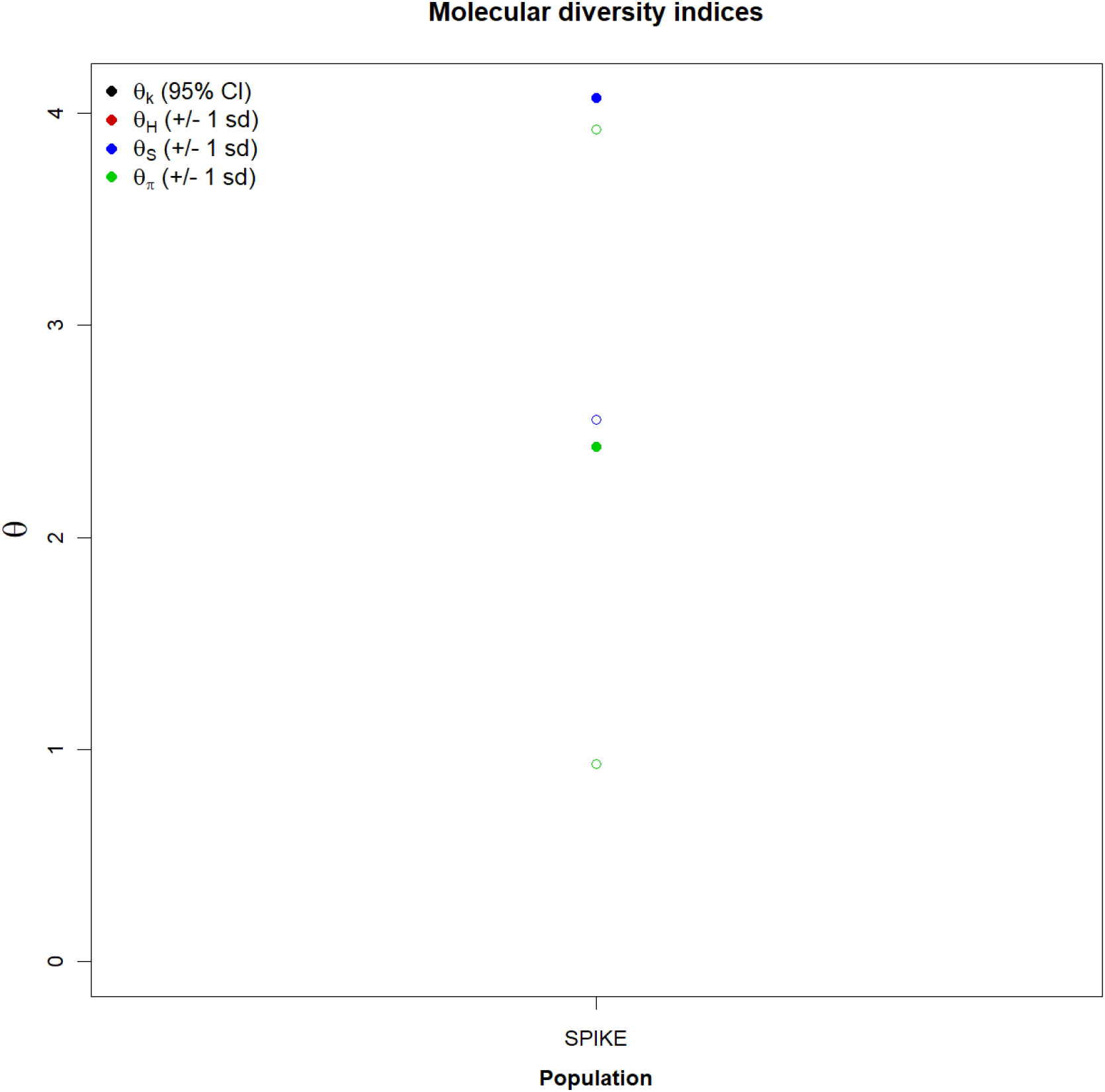
Graph of molecular diversity indices for the 37 haplotypes of the spike protein SARS-CoV-2. In the graph the values of θ: (θk) Relationship between the expected number of alllos (k) and the sample size; (θH) Expected homozygosity in a balanced relationship between drift and mutation; (θS) Relationship between the number of segregating sites (S), sample size (n) and non-recombinant sites; (θπ) Relationship between the average number of paired differences (π) and θ. * Generated by the statistical package in R language using the output data of the Arlequin software version 3.5.1.2.

**Figure 2.**
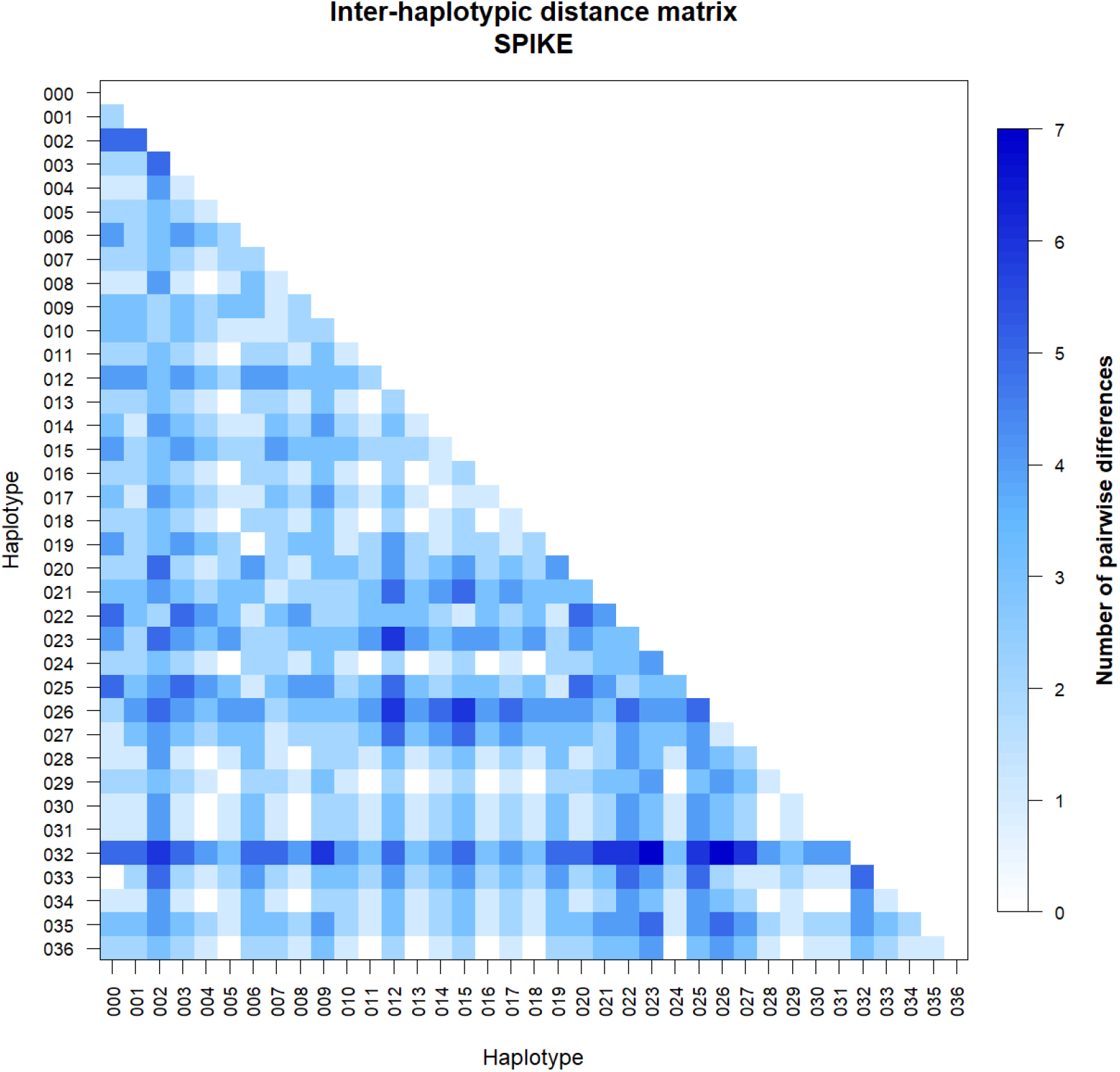
Inter haplotypic distance matrix for the 37 sequences of the spike protein SARS-CoV-2. *Generated by the statistical package in R language using the output data of the Software Arlequin version 3.5.1.2.

## 5. Discussion

With the use of methodologies for population structure analysis, it was possible to detect the existence of a high degree of similarity between haplotypes of the sars-cov-2 spike protein. Because significant levels of molecular diversity have not been found, we assume that molecular diversity for this protein, if found in future studies, may be associated with components of variation other than substitutions commonly found in the SARS-CoV-2 genome. These analyses support a consensus in the conservation of the Spike protein and it seems safe to state that the genetic variability of the virus, already found in some studies, is not mirrored in some of its genes, probably due to the high evolutionary degree of conservation of these genes, ensuring the expression of structural proteins.

All the considerations made in this study were supported by methodologies that ensured the discontinuous pattern of genetic divergence among haplotypes, taking into account a probable existence of many mutational stages. The values found for the genetic distance considered the minimum differences between the groups, as well as the inference of values greater than or equal to those observed in the proportion of these permutations, including the p-value of the test.

The φ estimators, although extremely sensitive to any form of molecular variation, supported the uniformity between the results found and all the methodologies employed, also clearly confirming the conservation of viral protein products. These considerations ensure that the use of neutralizing antibodies may be able to suppress the proliferation of the virus, further justifying the development of vaccines based on protein S.

